## Supplemental Information for "Integrative multi-omics profiling reveals cAMP-independent mechanisms regulating hyphal morphogenesis in *Candida albicans*"

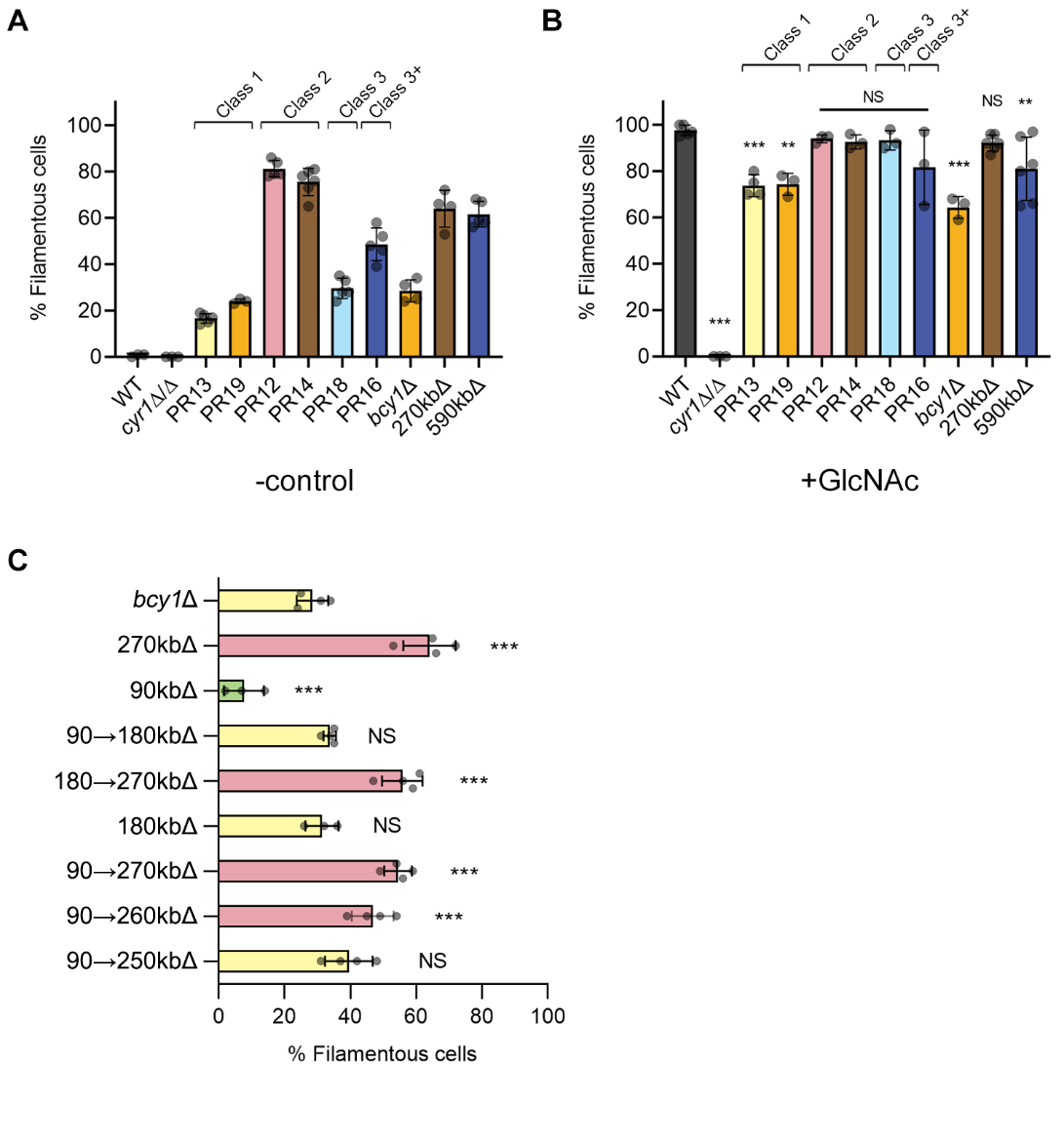

**S1 Fig. Hyphal induction rate of the fungal strains.**

(A) The plot shows the percent of filamentous cells in YPD medium at 30°C (-control).

(B) Graph indicating the percent of filamentous cells after growth in GlcNAc medium. Cells were grown in liquid medium containing 50 mM GlcNAc to induce hyphal growth at 37°C for 2 h and then filamentous cells were counted.

(C) The percent of filamentous cells in YPD medium at 30°C; green, weak filamentation; yellow, intermediate filamentation; pink, strong filamentation. Deletions indicated on the left are on chromosome 2 and they are heterozygous; the cells retain a wild-type version of chromosome 2.

(A, B, and C) Shown is the mean ± SD of at least 3 independent experiments with at least 100 cells counted for each condition. Statistical analysis was performed using one-way ANOVA with Dunnett's multiple comparisons test comparing the strains with the WT or parental strain; ^NS^ p > 0.05, ^**^ p < 0.01, ^***^ p < 0.001.

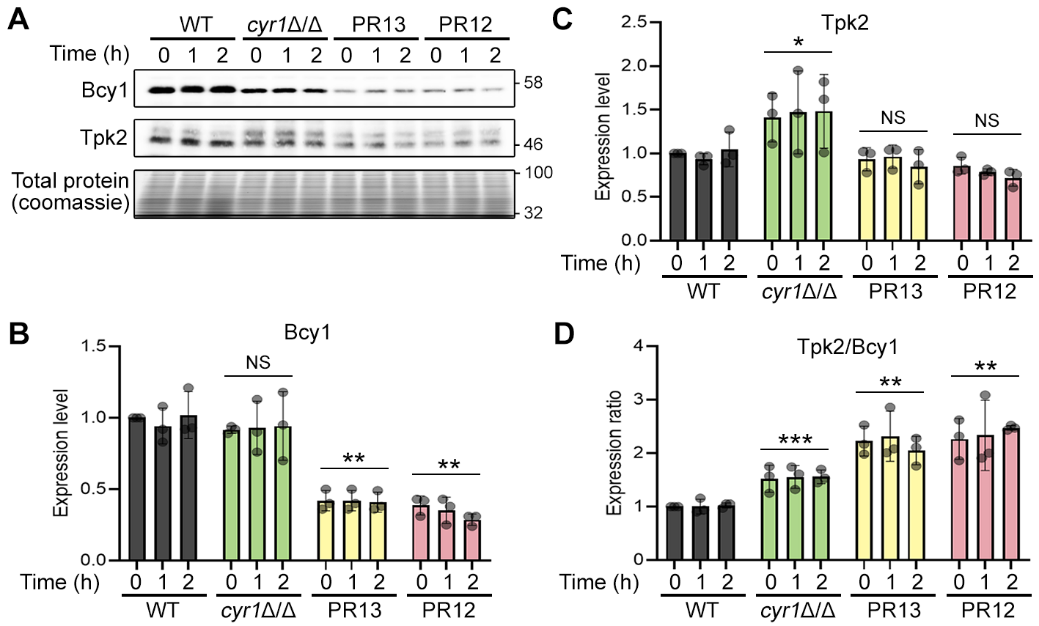

**S2 Fig. Expression levels of PKA subunits during hyphal induction.**

(A) Western blot detection of the negative regulatory subunit (Bcy1) and the catalytic subunit (Tpk2) of PKA. Cells were grown at 37°C in liquid galactose medium and then 50 mM GlcNAc was added for 2 h to induce hyphae. The sizes of the protein standards (kDa) are indicated on the right of each blot. Images shown are representative of three independent experiments.

(B-D) Relative levels of Bcy1 (B), Tpk2 (C), and Tpk2/Bcy1 ratio (D) compared to the WT 0-h samples. Shown is the mean ± SD of 3 independent experiments. Expression levels were normalized to total proteins on Coomassie-stained gels. Statistical analysis was performed using one-way ANOVA with Dunnett's multiple comparisons test comparing the strains with the WT; ^NS^ p > 0.05, ^*^ p < 0.05, ^**^ p < 0.01, ^***^ p < 0.001.

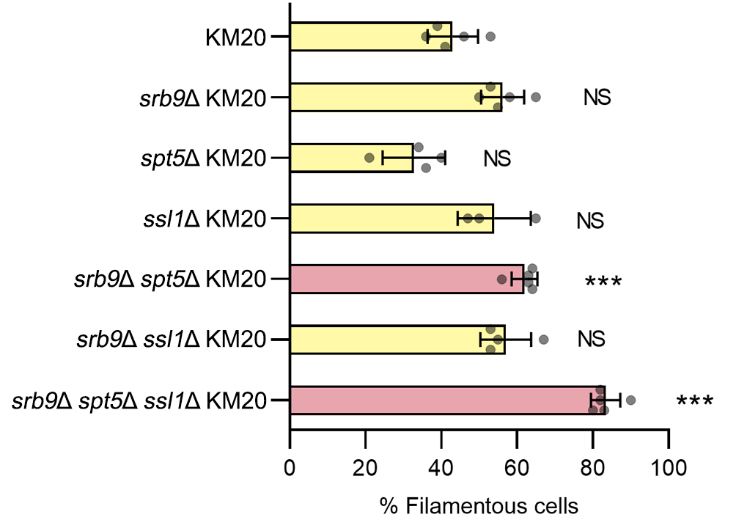

**S3 Fig. Gene-mapping analysis of the 10-kb region of chromosome 2 identified a role for *SRB9* and *SPT5* in hyphal induction in the PR mutants.**

The single, double, and triple heterozygous deletion mutants of *SRB9*, *SPT5*, and *SSL1* were created in KM20 strain background (*Chr2L 90kb→250kb∆ bcy1∆*/*BCY1 cyr1∆*/*∆*)*.* The plot shows the percent of filamentous cells in GlcNAc medium. Cells were grown in liquid medium containing 50 mM GlcNAc to induce hyphal growth at 37°C for 2 h and then filamentous cells were counted.; yellow, intermediate hyphal induction; pink, strong hyphal induction. Statistical analysis was performed using one-way ANOVA with Dunnett's multiple comparisons test comparing the strains with the parental strain; ^NS^ p > 0.01, ^***^ p < 0.001.

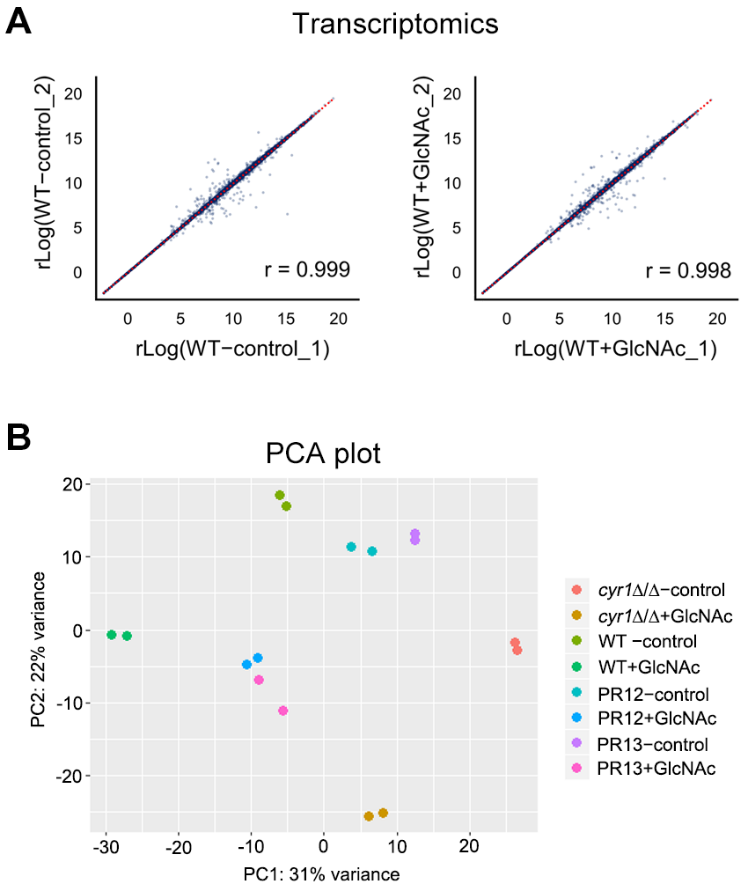

**S4 Fig. Correlation analysis and principal component analysis (PCA) of the normalized RNA-seq dataset.**

(A) Representative null comparisons of the biological replicates show very high reproducibility (r ≈ 1.00) in the RNA-seq dataset. Each dot represents the transcript level of individual gene in the scatter plots. We compared the biological replicates of WT−control and WT+GlcNAc. rLog, regularized log transformation.

(B) PCA plot shows clusters of biological replicates based on their similarity in transcriptome.

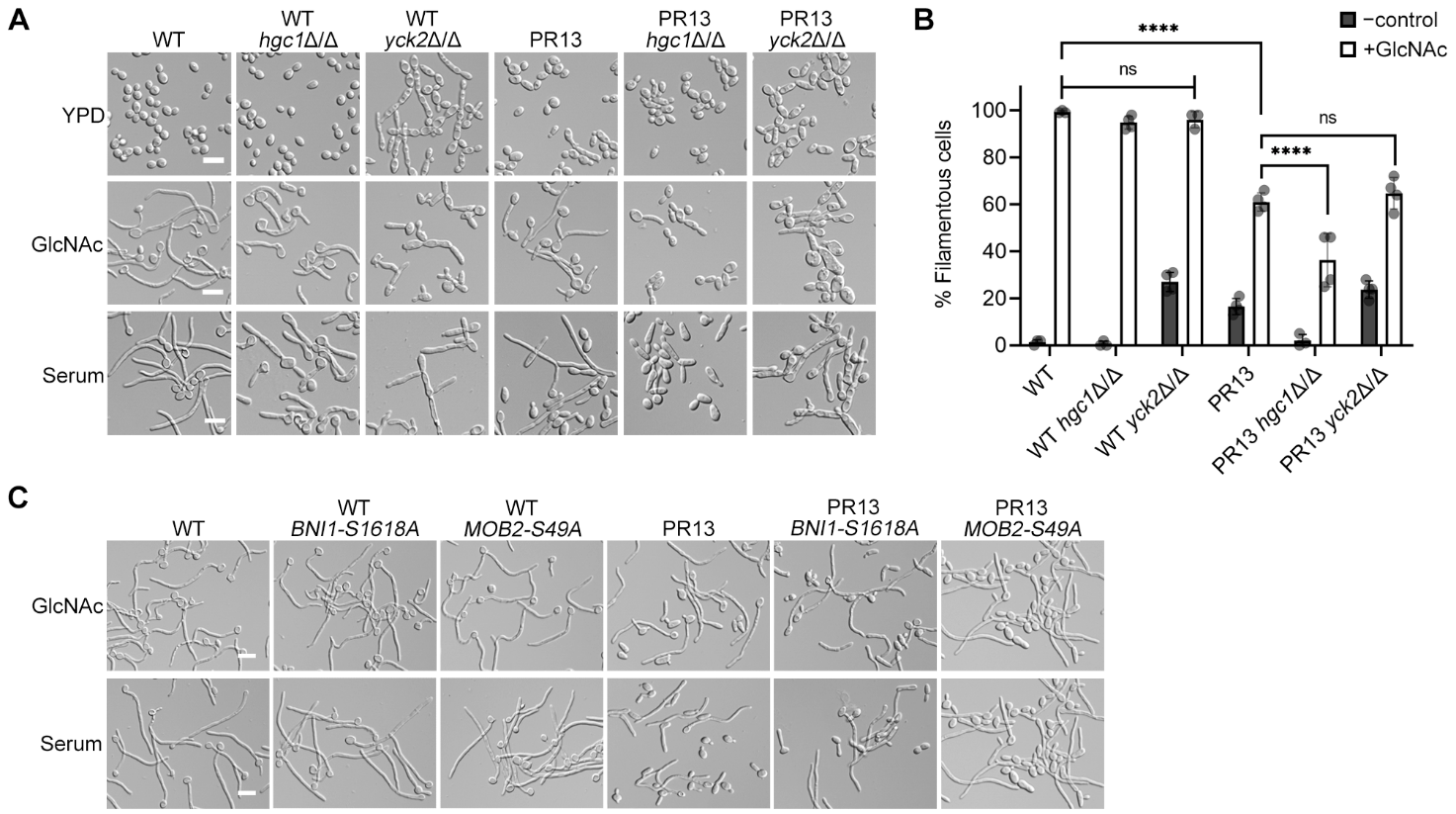

**S5 Fig. Deletion of Cdc28 cyclin (*HGC1*) and casein kinase 1 (*YCK2*) disrupt normal hyphal growth.**

(A) Deletion of Cdc28 cyclin (*HGC1*) and casein kinase 1 (*YCK2*) disrupt normal hyphal growth in WT and PR13 backgrounds.

(B) The plot shows the percent of filamentous cells in YPD medium at 30°C (-control) and after 2-h growth in GlcNAc medium at 37°C (+GlcNAc). Shown is the mean ± SD of at least 3 independent experiments with at least 100 cells counted for each condition. Statistical analysis was performed using one-way ANOVA with Dunnett's multiple comparisons test comparing the strains with the WT or parental strain; ^ns^ p > 0.05, ^****^ p < 0.0001.

(C) Phospho-mutants of Mob2 and Bni1 did not show an obvious defect in hyphal growth.

(A and C) The strains indicated at the top were grown in the liquid medium indicated on the left, and then hyphal induction was assessed microscopically. Cells were grown in liquid medium containing 15% serum or 50 mM N-acetylglucosamine (GlcNAc) to induce hyphal growth. Cells were incubated at 37°C for 2 h and then photographed. Scale bar, 10 μm.

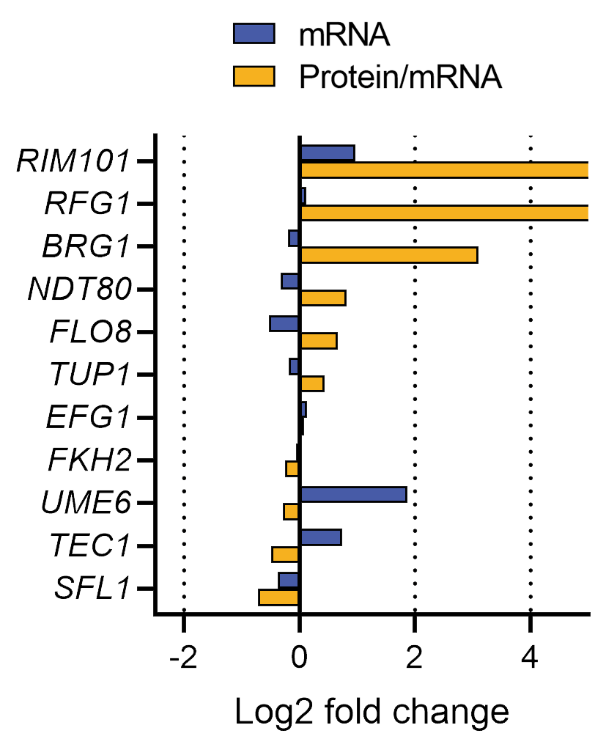

**S6 Fig.** **Protein-to-mRNA ratios of hyphal regulator TFs during GlcNAc induction in PR13.**

The relative change in protein-to-mRNA ratio for the selected 11 hyphal regulators TFs is shown in yellow, relative changes in mRNA expression are shown in blue. The protein-to-mRNA ratio of *RIM101* and *RFG1* increased dramatically (log2 fold change > 4) during hyphal induction while mRNA levels did not.

**S1 Table. Genome analysis summary**

| Strain | PR Class |  | Short genotype | *de novo* telomere addition^a^ | Telomere seed sequence |
| --- | --- | --- | --- | --- | --- |
| *cyr1*∆/∆ | - |  | *cyr1*∆/∆ | - | - |
| PR13 | 1 |  | *bcy1-Q82**/*BCY1 cyr1∆*/*∆* | - | - |
| PR19 | 1 |  | *bcy1-E19**/*BCY1 cyr1∆*/*∆* | - | - |
| PR12 | 2 |  | *Monosomy of ~276 kb of Chr2L cyr1∆*/*∆* | Detected at Chr2: 276,278 | ACATCCGTA |
| PR14 | 2 |  | *Monosomy of ~276 kb of Chr2L cyr1∆*/*∆* | Detected at Chr2: 276,278 | ACATCCGTA |
| PR18 | 3 |  | *Monosomy of ~590 kb of Chr2L cyr1∆*/*∆* | Detected at Chr2: 590,175 | CATCCG |
| PR16 | 3+ |  | *Monosomy of ~557 kb of Chr2L*  *Trisomy of ~1342 kb of Chr2 cyr1∆*/*∆* | Detected at Chr2: 1,899,373 | GAAG |
| PR2 | 3+ |  | *Monosomy of ~557 kb of Chr2L*  *Trisomy of ~1239 kb of Chr2 cyr1∆*/*∆* | Detected at Chr2: 1,796,423 | CGTACA |

^a^ 23-bp telomere repeat sequence, CACCAAGAAGTTAGACATCCGTA

**S2 Table. Potential phosphorylation substrates of Cdc28 and Yck2 during hyphal induction**

| Gene | Phosphosite | Description^a^ | Deletion phenotype^a^ |
| --- | --- | --- | --- |
| *BNI4* | S523 | Formin; Protein required for wild-type cell wall chitin distribution, morphology, hyphal growth | Hyphal induction: normal |
| *FPK1* | T143, S337 | Serine/threonine protein kinase | Hyphal induction: normal |
| *ZDS1* | S272 | Protein with a role in regulating Swe1p-dependent polarized growth | Hyphal induction: normal |
| *INT1* | S416, T1214 | Bud site selection protein Bud4 | Hyphal induction: normal |
| *SOL1* | T16 | Cell cycle regulator | Hyphal induction: normal |
| *BNI1* | S1619 | Formin; Role in cytoskeletal organization, cell polarity | Hyphal induction: abnormal |
| *MOB2* | S49 | Cbk1 kinase activator protein | Hyphal induction: absent |
| *ASK1* | S249 | Essential subunit of the Dam1 (DASH) complex, which acts in chromosome segregation by coupling kinetochores to spindle microtubules | Inviable |

^a^ Descriptions and deletion phenotypes were obtained from *Candida* Genome Database.

**S3 Table. *C. albicans* strains used in this study**

| **Strain** | **Short genotype** | **Parent or reference** | **Genotype** |
| --- | --- | --- | --- |
| DIC185 | Prototrophic wild type strain | (Wilson et al., 1999) | *ura3∆::λimm434*/*URA3 his1::hisG*/*HIS1 arg4::hisG*/*ARG4* |
| SP60-66 | *cyr1∆*/*∆* | (Parrino et al., 2017) | *cyr1∆::FRT*/*cyr1∆::ARG4 ura3∆::λimm434*/*URA3 his1::hisG*/*HIS1 arg4::hisG*/*arg4::hisG* |
| PR2 | *Monosomy of ~557 kb of Chr2L*  *Trisomy of ~1239 kb of Chr2*  *cyr1∆*/*∆* | SP60-66 | *Monosomy of ~557 kb of Chr2L (Chr2L→C2_02790C)*  *Trisomy of ~1239 kb of Chr2 (C2_02800W→C2_08860W)*  *cyr1∆::FRT*/*cyr1∆::ARG4 ura3∆::λimm434*/*URA3 his1::hisG*/*HIS1 arg4::hisG*/*arg4::hisG* |
| PR12 | *Monosomy of ~276 kb of Chr2L*  *cyr1∆*/*∆* | SP60-66 | *Monosomy of ~276 kb of Chr2L (Chr2L→C2_01540W)*  *cyr1∆::FRT*/*cyr1∆::ARG4 ura3∆::λimm434*/*URA3 his1::hisG*/*HIS1 arg4::hisG*/*arg4::hisG* |
| PR13 | *bcy1-Q82**/*BCY1 cyr1∆*/*∆* | SP60-66 | *bcy1-Q82**/*BCY1 cyr1∆::FRT*/*cyr1∆::ARG4 ura3∆::λimm434*/*URA3 his1::hisG*/*HIS1 arg4::hisG*/*arg4::hisG* |
| PR14 | *Monosomy of ~276 kb of Chr2L*  *cyr1∆*/*∆* | SP60-66 | *Monosomy of ~276 kb of Chr2L (Chr2L→C2_01540W)*  *cyr1∆::FRT*/*cyr1∆::ARG4 ura3∆::λimm434*/*URA3 his1::hisG*/*HIS1 arg4::hisG*/*arg4::hisG* |
| PR16 | *Monosomy of ~557 kb of Chr2L*  *Trisomy of ~1342 kb of Chr2*  *cyr1∆*/*∆* | SP60-66 | *Monosomy of ~557 kb of Chr2L (Chr2L→C2_02790C)*  *Trisomy of ~1342 kb of Chr2 (C2_02800W→C2_09290W)*  *cyr1∆::FRT*/*cyr1∆::ARG4 ura3∆::λimm434*/*URA3 his1::hisG*/*HIS1 arg4::hisG*/*arg4::hisG* |
| PR18 | *Monosomy of ~590 kb of Chr2L*  *cyr1∆*/*∆* | SP60-66 | *Monosomy of ~590 kb of Chr2L (Chr2L→C2_02960C)*  *cyr1∆::FRT*/*cyr1∆::ARG4 ura3∆::λimm434*/*URA3 his1::hisG*/*HIS1 arg4::hisG*/*arg4::hisG* |
| PR19 | *bcy1-E19**/*BCY1 cyr1∆*/*∆* | SP60-66 | *bcy1-E19**/*BCY1 cyr1∆::FRT*/*cyr1∆::ARG4 ura3∆::λimm434*/*URA3 his1::hisG*/*HIS1 arg4::hisG*/*arg4::hisG* |
| KM11 | *bcy1∆*/*BCY1 cyr1∆*/*∆* | SP60-66 | *bcy1∆::SAT1-FLIP*/*BCY1 cyr1∆::FRT*/*cyr1∆::ARG4 ura3∆::λimm434*/*URA3 his1::hisG*/*HIS1 arg4::hisG*/*arg4::hisG* |
| KM12 | *Monosomic:C2_00030W→C2_01540W*  *(Chr2L 270kb deletion)*  *bcy1∆*/*BCY1 cyr1∆*/*∆* | SP60-66 | *c2_00030w→c2_01540wΔ::SAT1-FLIP*/*C2_00030W→C2_01540W cyr1∆::FRT*/*cyr1∆::ARG4 ura3∆::λimm434*/*URA3 his1::hisG*/*HIS1 arg4::hisG*/*arg4::hisG* |
| KM13 | *Monosomic:C2_00030W→C2_02960C*  *(Chr2L 590kb deletion)*  *bcy1∆*/*BCY1 cyr1∆*/*∆* | SP60-66 | *c2_00030w→c2_02960cΔ::SAT1-FLIP*/*C2_00030W→C2_02960C cyr1∆::FRT*/*cyr1∆::ARG4 ura3∆::λimm434*/*URA3 his1::hisG*/*HIS1 arg4::hisG*/*arg4::hisG* |
| KM14 | *Monosomic:C2_00030W→C2_00550W*  *(Chr2L 90kb deletion)*  *bcy1∆*/*BCY1 cyr1∆*/*∆* | KM11 | *c2_00030w→c2_00550wΔ::SAT1-FLIP*/*C2_00030W→C2_00550W cyr1∆::FRT*/*cyr1∆::ARG4 ura3∆::λimm434*/*URA3 his1::hisG*/*HIS1 arg4::hisG*/*arg4::hisG* |
| KM15 | *Monosomic:C2_00560W→C2_01140C*  *(Chr2L 90kb→180kb deletion)*  *bcy1∆*/*BCY1 cyr1∆*/*∆* | KM11 | *c2_00560w→c2_01140cΔ::SAT1-FLIP*/*C2_00560W→C2_01140C cyr1∆::FRT*/*cyr1∆::ARG4 ura3∆::λimm434*/*URA3 his1::hisG*/*HIS1 arg4::hisG*/*arg4::hisG* |
| KM16 | *Monosomic:C2_01150W→C2_01540W*  *(Chr2L 180kb→270kb deletion)*  *bcy1∆*/*BCY1 cyr1∆*/*∆* | KM11 | *c2_01150w→c2_01540wΔ::SAT1-FLIP*/*C2_01150W→C2_01540W cyr1∆::FRT*/*cyr1∆::ARG4 ura3∆::λimm434*/*URA3 his1::hisG*/*HIS1 arg4::hisG*/*arg4::hisG* |
| KM17 | *Monosomic:C2_00030W→C2_01140C*  *(Chr2L 180kb deletion)*  *bcy1∆*/*BCY1 cyr1∆*/*∆* | KM11 | *c2_00030w→c2_01140cΔ::SAT1-FLIP*/*C2_00030W→C2_01140C cyr1∆::FRT*/*cyr1∆::ARG4 ura3∆::λimm434*/*URA3 his1::hisG*/*HIS1 arg4::hisG*/*arg4::hisG* |
| KM18 | *Monosomic:C2_00560W→C2_01540W*  *(Chr2L 90kb→270kb deletion)*  *bcy1∆*/*BCY1 cyr1∆*/*∆* | KM11 | *c2_00560w→c2_01540wΔ::SAT1-FLIP*/*C2_00560W→C2_01540W cyr1∆::FRT*/*cyr1∆::ARG4 ura3∆::λimm434*/*URA3 his1::hisG*/*HIS1 arg4::hisG*/*arg4::hisG* |
| KM19 | *Monosomic:C2_00560W→C2_01500W*  *(Chr2L 90kb→260kb deletion)*  *bcy1∆*/*BCY1 cyr1∆*/*∆* | KM11 | *c2_00560w→c2_01500wΔ::SAT1-FLIP*/*C2_00560W→C2_01500W cyr1∆::FRT*/*cyr1∆::ARG4 ura3∆::λimm434*/*URA3 his1::hisG*/*HIS1 arg4::hisG*/*arg4::hisG* |
| KM20 | *Monosomic:C2_00560W→C2_01460C*  *(Chr2L 90kb→250kb deletion)*  *bcy1∆*/*BCY1 cyr1∆*/*∆* | KM11 | *c2_00560w→c2_01460cΔ::SAT1-FLIP*/*C2_00560W→C2_01460C cyr1∆::FRT*/*cyr1∆::ARG4 ura3∆::λimm434*/*URA3 his1::hisG*/*HIS1 arg4::hisG*/*arg4::hisG* |
| KM21 | *hgc1∆*/*∆* | DIC185 | *hgc1∆::SAT1-FLIP*/*hgc1∆::SAT1-FLIP ura3∆::λimm434*/*URA3 his1::hisG*/*HIS1 arg4::hisG*/*ARG4* |
| KM22 | *yck2∆*/*∆* | DIC185 | *yck2∆::SAT1-FLIP*/*yck2∆::SAT1-FLIP ura3∆::λimm434*/*URA3 his1::hisG*/*HIS1 arg4::hisG*/*ARG4* |
| KM23 | *hgc1∆*/*∆* PR13 | PR13 | *hgc1∆::SAT1-FLIP*/*hgc1∆::SAT1-FLIP bcy1-Q82**/*BCY1 cyr1∆::FRT*/*cyr1∆::ARG4 ura3∆::λimm434*/*URA3 his1::hisG*/*HIS1 arg4::hisG*/*arg4::hisG* |
| KM24 | *yck2∆*/*∆* PR13 | PR13 | *yck2∆::SAT1-FLIP*/*yck2∆::SAT1-FLIP bcy1-Q82**/*BCY1 cyr1∆::FRT*/*cyr1∆::ARG4 ura3∆::λimm434*/*URA3 his1::hisG*/*HIS1 arg4::hisG*/*arg4::hisG* |
| KM25 | *BNI1-S1618A* | DIC185 | *BNI1-S1618A*/*BNI1-S1618A eno1::CaCAS9-SAT1-sgRNA*/*ENO1 ura3∆::λimm434*/*URA3 his1::hisG*/*HIS1 arg4::hisG*/*ARG4* |
| KM26 | *MOB2-S49A* | DIC185 | *MOB2-S49A*/*MOB2-S49A eno1::CaCAS9-SAT1-sgRNA*/*ENO1 ura3∆::λimm434*/*URA3 his1::hisG*/*HIS1 arg4::hisG*/*ARG4* |
| KM27 | *BNI1-S1618A* PR13 | PR13 | *BNI1-S1618A*/*BNI1-S1618A eno1::CaCAS9-SAT1-sgRNA*/*ENO1 bcy1-Q82**/*BCY1 cyr1∆::FRT*/*cyr1∆::ARG4 ura3∆::λimm434*/*URA3 his1::hisG*/*HIS1 arg4::hisG*/*arg4::hisG* |
| KM28 | *MOB2-S49A* PR13 | PR13 | *MOB2-S49A*/*MOB2-S49A eno1::CaCAS9-SAT1-sgRNA*/*ENO1 bcy1-Q82**/*BCY1 cyr1∆::FRT*/*cyr1∆::ARG4 ura3∆::λimm434*/*URA3 his1::hisG*/*HIS1 arg4::hisG*/*arg4::hisG* |
| KM29 | *srb9∆* KM20 | KM20 | *srb9∆::SAT1-FLIP*/*SRB9* c2*_00560w→c2_01460cΔ::SAT1-FLIP*/*C2_00560W→C2_01460C*  *cyr1∆::FRT*/*cyr1∆::ARG4 ura3∆::λimm434*/*URA3 his1::hisG*/*HIS1 arg4::hisG*/*arg4::hisG* |
| KM30 | *spt5∆* KM20 | KM20 | *spt5∆::SAT1-FLIP*/*SPT5 c2_00560w→c2_01460cΔ::SAT1-FLIP*/*C2_00560W→C2_01460C*  *cyr1∆::FRT*/*cyr1∆::ARG4 ura3∆::λimm434*/*URA3 his1::hisG*/*HIS1 arg4::hisG*/*arg4::hisG* |
| KM31 | *ssl1∆* KM20 | KM20 | *ssl1∆::SAT1-FLIP*/*SSL1 c2_00560w→c2_01460cΔ::SAT1-FLIP*/*C2_00560W→C2_01460C*  *cyr1∆::FRT*/*cyr1∆::ARG4 ura3∆::λimm434*/*URA3 his1::hisG*/*HIS1 arg4::hisG*/*arg4::hisG* |
| KM32 | *srb9∆* *spt5∆* KM20 | KM20 | *srb9∆::SAT1-FLIP*/*SRB9* *spt5∆::SAT1-FLIP*/*SPT5 c2_00560w→c2_01460cΔ::SAT1-FLIP*/*C2_00560W→C2_01460C cyr1∆::FRT*/*cyr1∆::ARG4 ura3∆::λimm434*/*URA3 his1::hisG*/*HIS1 arg4::hisG*/*arg4::hisG* |
| KM33 | *srb9∆* *ssl1∆* KM20 | KM20 | *srb9∆::SAT1-FLIP*/*SRB9* *ssl1∆::SAT1-FLIP*/*SSL1 c2_00560w→c2_01460cΔ::SAT1-FLIP*/*C2_00560W→C2_01460C cyr1∆::FRT*/*cyr1∆::ARG4 ura3∆::λimm434*/*URA3 his1::hisG*/*HIS1 arg4::hisG*/*arg4::hisG* |

**S4 Table. Target sequences of the sgRNAs**

| **Target gene** | **20-nt target sequence** |
| --- | --- |
| *C2_00030W* | TCAAGACGATCTGAAATTGG |
| *C2_01540W* | GAGCCAAAAGATACAAGCGG |
| *C2_02960C* | TTGTAGCCAAGTCATACCCG |
| *C2_00550W* | TCATCAAAAAGAGCCAGTGT |
| *C2_00560W* | GATGCTATCGAGGAAGAAGT |
| *C2_01140C* | TTCAACATAGAAGTCCATAT |
| *C2_01150W* | TTTAATTGTAACAAGTACCG |
| *C2_01300C* | ATGATGATAAAGTTGACCGT |
| *C2_01310W* | ATGGGTGGGATTATTAATGT |
| *C2_01430W* | AAAGCTTTCAAATTACACGT |
| *C2_01420C* | AACAGATATAGTCTATGCTA |
| *C2_01460C* | GTTAAATATTAAAAAGTGCT |
| *C2_01500W* | TGTATTTCAATATATGCCAA |
| *C2_00770W* | AGATAAAGCTTTAGACATTG |
| *C2_01000W* | CTCATCAACATCTACAGCTG |
| *HGC1* | GTAGTACTACATGATGAACT |
| *YCK2* | CCAGCAGTTACATTATGTGA |

**S5 Table. The oligonucleotides used in this study**

| **Name** | **Sequence (5' to 3')** | **Description** |
| --- | --- | --- |
| SNR52_R_30 | ccaatttcagatcgtcttgaCAAATTAAAAATAGTTTACGCAAGTC | sgRNA synthesis against *C2_00030W* |
| sgRNA_F_30 | tcaagacgatctgaaattggGTTTTAGAGCTAGAAATAGCAAGTTAAA | sgRNA synthesis against *C2_00030W* |
| SNR52_R_1540 | ccgcttgtatcttttggctcCAAATTAAAAATAGTTTACGCAAGTC | sgRNA synthesis against *C2_01540W* |
| sgRNA_F_1540 | gagccaaaagatacaagcggGTTTTAGAGCTAGAAATAGCAAGTTAAA | sgRNA synthesis against *C2_01540W* |
| SNR52_R_2960 | cgggtatgacttggctacaaCAAATTAAAAATAGTTTACGCAAGTC | sgRNA synthesis against *C2_02960C* |
| sgRNA_F_2960 | ttgtagccaagtcatacccgGTTTTAGAGCTAGAAATAGCAAGTTAAA | sgRNA synthesis against *C2_02960C* |
| 30_NAT_FLP_For | ctagaaaccgatgccttgcccatcaacaccatcacagataaaatcaatctccccattggccaccccgaatcaatcacactTAAAGGGAACAAAAGCTGGG | Deletion construct with a homology arm to *C2_00030W* |
| 1540_NAT_FLP_Rev | cttgtggttgagatttcacagagcaaccatctttcttcaaaacagactgatcatgtctaaactctttaccatgaccataaCTCTAGAACTAGTGGATCTG | Deletion construct with a homology arm to *C2_01540W* |
| 2960_NAT_FLP_Rev | tggatctctagaacattttgaaaatacagcaatagacccggataccatttttcaggagaagggtgtaattgaattgaacaCTCTAGAACTAGTGGATCTG | Deletion construct with a homology arm to *C2_02960C* |
| Chr2Del_F1 | TACCGGTCTTCCCATCAACT | Genotype PCR forward primer binding at Chr2A: 5,510..5,529 |
| Chr2Del_F2 | TCAGCAAAATGATCGAGGCA | Genotype PCR forward primer binding at Chr2A: 274,792..274,811 |
| Chr2Del_R1 | AATGGAGCGTCAGCTAAAGG | Genotype PCR reverse primer binding at Chr2A: 275,578..275,559 |
| Chr2Del_R2 | AGCGATTGCTGGTCTCAAAA | Genotype PCR reverse primer binding at Chr2A: 276,530..276,511 |
| Chr2Del_R3 | CACCATGACATTCGGTGCTA | Genotype PCR reverse primer binding at Chr2A: 591,190..591,171 |
| SNR52_R_550 | acactggctctttttgatgaCAAATTAAAAATAGTTTACGCAAGTC | sgRNA synthesis against *C2_00550W* |
| sgRNA_F_550 | tcatcaaaaagagccagtgtGTTTTAGAGCTAGAAATAGCAAGTTAAA | sgRNA synthesis against *C2_00550W* |
| 550_NAT_FLP_Rev | cagccaccgccaaagcatccaagttagttgtcacatgattggtgactctatttgatagaatattgtgtatttcacctttaCTCTAGAACTAGTGGATCTG | Deletion construct with a homology arm to *C2_00550W* |
| SNR52_R_560 | acttcttcctcgatagcatcCAAATTAAAAATAGTTTACGCAAGTC | sgRNA synthesis against *C2_00560W* |
| sgRNA_F_560 | gatgctatcgaggaagaagtGTTTTAGAGCTAGAAATAGCAAGTTAAA | sgRNA synthesis against *C2_00560W* |
| 560_NAT_FLP_For | aaactattacgttaaatctgataatgaacaattcaaatgattggagtttttcattatgtgatgactctcttgaactatttTAAAGGGAACAAAAGCTGGG | Deletion construct with a homology arm to *C2_00560W* |
| SNR52_R_1140 | atatggacttctatgttgaaCAAATTAAAAATAGTTTACGCAAGTC | sgRNA synthesis against *C2_01140C* |
| sgRNA_F_1140 | ttcaacatagaagtccatatGTTTTAGAGCTAGAAATAGCAAGTTAAA | sgRNA synthesis against *C2_01140C* |
| 1140_NAT_FLP_Rev | ttgtattcatattgagtcccatcttttgcaaccaacacaggcatattcttgtacgtcagtgaattagggatgaaagggacCTCTAGAACTAGTGGATCTG | Deletion construct with a homology arm to *C2_01140C* |
| SNR52_R_1150 | cggtacttgttacaattaaaCAAATTAAAAATAGTTTACGCAAGTC | sgRNA synthesis against *C2_01150W* |
| sgRNA_F_1150 | tttaattgtaacaagtaccgGTTTTAGAGCTAGAAATAGCAAGTTAAA | sgRNA synthesis against *C2_01150W* |
| 1150_NAT_FLP_For | agctaatatctgcaaatcaattagagtcattattatctccatcattacgatacattttggttcactatgccagtaaatatTAAAGGGAACAAAAGCTGGG | Deletion construct with a homology arm to *C2_01150W* |
| Chr2Del_F3 | CGAACTTGACGACCTAGACG | Genotype PCR forward primer binding at Chr2A: 6,351..6,370 |
| Chr2Del_R4 | TCGTCACAGAACTTTAGCGG | Genotype PCR reverse primer binding at Chr2A: 7,055..7,036 |
| Chr2Del_R5 | AAGAGGACCAACCAGTACCA | Genotype PCR reverse primer binding at Chr2A: 88,739..88,720 |
| Chr2Del_F4 | GCCCTGCAAGTATCCAATCA | Genotype PCR forward primer binding at Chr2A: 89,570..89,589 |
| Chr2Del_F5 | CAACCTCTCCCCAAACTCAC | Genotype PCR forward primer binding at Chr2A: 90,395..90,414 |
| Chr2Del_R7 | GAGGTTCTGGTTCTTCGGTG | Genotype PCR reverse primer binding at Chr2A: 91,229..91,210 |
| Chr2Del_R6 | TTGGGGTTGTTTTCCGACAT | Genotype PCR reverse primer binding at Chr2A: 185,045..185,064 |
| Chr2Del_F6 | AACTCAGTCATTGGACACGC | Genotype PCR forward primer binding at Chr2A: 186,488..186,507 |
| 1140_NAT_FLP_Rev | atcaatgcaaagcttgaaactgattattgaaactaataaaatcaggccagcaaaacttaaggagtttgatttgatatttaCTCTAGAACTAGTGGATCTG | Deletion construct with a homology arm to *C2_01140C* |
| Chr2Del_R8 | TTCTGTGAATGTCCCTTGCC | Genotype PCR reverse primer binding at Chr2A: 186887..186868 |
| 1300_NAT_FLP_Rev | atcccagtatcatcaaaaactgttcgtcttatacttgtgtcattacttctaataacactaataaatatattggctgctttCTCTAGAACTAGTGGATCTG | Deletion construct with a homology arm to *C2_01300C* |
| SNR52_R_1300 | acggtcaactttatcatcatCAAATTAAAAATAGTTTACGCAAGTC | sgRNA synthesis against *C2_01300C* |
| sgRNA_F_1300 | atgatgataaagttgaccgtGTTTTAGAGCTAGAAATAGCAAGTTAAA | sgRNA synthesis against *C2_01300C* |
| 1310_NAT_FLP_For | catcaccaacaacgaaaatagttgttaaaagacgatatagtgaatttaaatctttgagagacaatttattaaaattattcTAAAGGGAACAAAAGCTGGG | Deletion construct with a homology arm to *C2_01310W* |
| SNR52_R_1310 | acattaataatcccacccatCAAATTAAAAATAGTTTACGCAAGTC | sgRNA synthesis against *C2_01310W* |
| sgRNA_F_1310 | atgggtgggattattaatgtGTTTTAGAGCTAGAAATAGCAAGTTAAA | sgRNA synthesis against *C2_01310W* |
| 1420_NAT_FLP_Rev | agtatgatcagaatattagtgattcagaacacgatttaacaccaatcaaaagaaagcgtcaatcagcacaatcggcaccaCTCTAGAACTAGTGGATCTG | Deletion construct with a homology arm to *C2_01420C* |
| SNR52_R_1420 | ccgccagcaacaaagttttcCAAATTAAAAATAGTTTACGCAAGTC | sgRNA synthesis against *C2_01420C* |
| sgRNA_F_1420 | gaaaactttgttgctggcggGTTTTAGAGCTAGAAATAGCAAGTTAAA | sgRNA synthesis against *C2_01420C* |
| 1430_NAT_FLP_For | ctaacttcccaaaagaaggaattttatttgaagatttcttaccaattttcactaagccagacttgtttaataaattagtcTAAAGGGAACAAAAGCTGGG | Deletion construct with a homology arm to *C2_01430W* |
| SNR52_R_1430 | acgtgtaatttgaaagctttCAAATTAAAAATAGTTTACGCAAGTC | sgRNA synthesis against *C2_01430W* |
| sgRNA_F_1430 | aaagctttcaaattacacgtGTTTTAGAGCTAGAAATAGCAAGTTAAA | sgRNA synthesis against *C2_01430W* |
| Chr2Del_R9 | TTGCTCGTCCCCTTTCATAC | Genotype PCR reverse primer binding at Chr2A: 221,879..221,860 |
| SNR52_R_1420_new | TAGCATAGACTATATCTGTTCAAATTAAAAATAGTTTACGCAAGTC | sgRNA synthesis against *C2_01420C* |
| sgRNA_F_1420_new | AACAGATATAGTCTATGCTAGTTTTAGAGCTAGAAATAGCAAGTTAAA | sgRNA synthesis against *C2_01420C* |
| 1420_NAT_Rev_new | AAGTTTTACTCTTGAACAATCACCAGAAATAAAACCTAAACCTAAATCAAAAACTTCAGATTTAACAGATATAGTCTATGCTCTAGAACTAGTGGATCTG | Deletion construct with a homology arm to *C2_01420C* |
| Chr2Del_R10 | GTACAGACCGACACAACTCC | Genotype PCR reverse primer binding at Chr2A: 256,183..256,164 |
| SNR52_R_1460 | AGCACTTTTTAATATTTAACCAAATTAAAAATAGTTTACGCAAGTC | sgRNA synthesis against *C2_01460C* |
| sgRNA_F_1460 | GTTAAATATTAAAAAGTGCTGTTTTAGAGCTAGAAATAGCAAGTTAAA | sgRNA synthesis against *C2_01460C* |
| 1460_NAT_FLP_Rev | GCAACACTTTAACAAAAAGTGTATTCATGTTCTATAAGAACAATTGAGAAGACAGAGTATAAACCCAACATTGCTTACAACTCTAGAACTAGTGGATCTG | Deletion construct with a homology arm to *C2_01460C* |
| Chr2Del_R11 | AACCTAGCATTGATGGAGCC | Genotype PCR reverse primer binding at Chr2A: 260,529..260,510 |
| SNR52_R_1500 | TTGGCATATATTGAAATACACAAATTAAAAATAGTTTACGCAAGTC | sgRNA synthesis against *C2_01500W* |
| sgRNA_F_1500 | TGTATTTCAATATATGCCAAGTTTTAGAGCTAGAAATAGCAAGTTAAA | sgRNA synthesis against *C2_01500W* |
| 1500_NAT_FLP_Rev | TTCTGGTGAGTACGTAGAGAGTAATTGAGCAACAAGATCCTAAAAAAAAAAGCGTAGTCTAGTAGAGAAAAACAATTCAGCTCTAGAACTAGTGGATCTG | Deletion construct with a homology arm to *C2_01500W* |
| Chr2Del_R12 | ACAAAATGAGTCGGCACCAA | Genotype PCR reverse primer binding at Chr2A: 271,047..271,028 |
| 1470_NAT_FLP_For | TTTAACTTCAGCTTTCTTCTGGCTACTACAATCTTCACTTACAACTAGAATTCATACTAAAAACATTCGATAGAAACATCTAAAGGGAACAAAAGCTGGG | Deletion construct with a homology arm to *C2_01470W* |
| 1470_NAT_FLP_Rev | AAGTCGTGCGAAATGGCCATACTCATTAAACCTATGTACAGCTTTCAATCTTTATCTACATATATGTGCACTAAATAATCCTCTAGAACTAGTGGATCTG | Deletion construct with a homology arm to *C2_01470W* |
| 1480_NAT_FLP_For | AACAGAGATCTCTGGTCATTAATATTCACAGCAATTTACACATCATCTCAACTATATCAAATGTCAGACAGTGATCTACCTAAAGGGAACAAAAGCTGGG | Deletion construct with a homology arm to *C2_01480W* |
| 1480_NAT_FLP_Rev | ATCAACATAAATAAACAAAGACAGTAGTGAAGAACTAACAAGGGAGCGCACTTCAGTTAGAAAAACAACATTTCTTGGTACTCTAGAACTAGTGGATCTG | Deletion construct with a homology arm to *C2_01480W* |
| 1490_NAT_FLP_For | AATACCAAATATAATTTTAATTGTTTTCACAACCTGGACAATTATGAAGAACTTCGTGCACAAAAACATCACAGTTGATATAAAGGGAACAAAAGCTGGG | Deletion construct with a homology arm to *C2_01490C* |
| 1490_NAT_FLP_Rev | TGGGCTCTAAATCGCCGTCCTCTAAACTTCCTTCGACTGCTCCAGAGAAAAGTGCATCACCTGGAGGAGCAAAGAGTGGTCTCTAGAACTAGTGGATCTG | Deletion construct with a homology arm to *C2_01490C* |
| SNR52_R_770 | CAATGTCTAAAGCTTTATCTCAAATTAAAAATAGTTTACGCAAGTC | sgRNA synthesis against *C2_00770W* |
| sgRNA_F_770 | AGATAAAGCTTTAGACATTGGTTTTAGAGCTAGAAATAGCAAGTTAAA | sgRNA synthesis against *C2_00770W* |
| Chr2Del_F7 | GAGAGTTAACGGCGTGACTT | Genotype PCR forward primer binding at Chr2A: 131,649..131,668 |
| 770_NAT_FLP_For | ACTACACCACGCTACACATTTTCTTTTATACCCCTTGAATTAACAATTCTATTTCAAACAAATACTTATAATCTTCATAATAAAGGGAACAAAAGCTGGG | Deletion construct with a homology arm to *C2_00770W* |
| SNR52_R_1000 | CAGCTGTAGATGTTGATGAGCAAATTAAAAATAGTTTACGCAAGTC | sgRNA synthesis against *C2_01000W* |
| sgRNA_F_1000 | CTCATCAACATCTACAGCTGGTTTTAGAGCTAGAAATAGCAAGTTAAA | sgRNA synthesis against *C2_01000W* |
| Chr2Del_F8 | ACACAATCACCCTACACTGC | Genotype PCR forward primer binding at Chr2A: 163,901..163,920 |
| 1000_NAT_FLP_For | CCGAGTTTAGTTTCATTTTTCCTTTTTCTCTTTTTCTACATCATCCTCACAACAATTTCAAATATGTCTCAAGACAACGTTAAAGGGAACAAAAGCTGGG | Deletion construct with a homology arm to *C2_01000W* |
| 1470_check_For | AGTGAAGTAGGAGACACACGA | Genotype PCR primer for detecting *C2_01470W* allele |
| 1470_check_Rev | TCGACCTTGTTGTGACCAAA | Genotype PCR primer for detecting *C2_01470W* allele |
| 1480_check_For | GGTATTGACCCTGAGGTGGT | Genotype PCR primer for detecting *C2_01480W* allele |
| 1480_check_Rev | TTGATCACCGGCTCTTCTGA | Genotype PCR primer for detecting *C2_01480W* allele |
| 1490_check_For | CTGCTTGGAACGGAGGTATG | Genotype PCR primer for detecting *C2_01490C* allele |
| 1490_check_Rev | AGCCCAATATACGACAGCAC | Genotype PCR primer for detecting *C2_01490C* allele |
| BCY1_For | TGTGATACCGGTCTTTTAGCA | Genotype PCR primer for detecting *BCY1* allele |
| BCY1_Rev | CACCAGAAGGTGAGTAAGCC | Genotype PCR primer for detecting *BCY1* allele |
| BCY1_NAT_FLP_For | ATGTTGCCCACAAATAAATTGTGATACCGGTCTTTTAGCATATATCTTCTACTCTTCAATCAACATCTTTACCAATGTCTTAAAGGGAACAAAAGCTGGG | *BCY1* deletion construct |
| BCY1_NAT_FLP_Rev | GGTGAGTAAGCCGAAAAAAAAAAGATAGAATATACTAACATAAAACAATATAAGATAATAAACAACTACTTATTGTACACCTCTAGAACTAGTGGATCTG | *BCY1* deletion construct |
| sgMOB2_top | atttgTTTTGAAGATAATTTTGAAGg | CRISPR plasmid cloning against *MOB2* |
| sgMOB2_bottom | aaaacCTTCAAAATTATCTTCAAAAc | CRISPR plasmid cloning against *MOB2* |
| MOB2_S49A_For | ATCTGGCAATGGTTTGAGACGAACACAATCACCTACCAAGTTTgCgCCaTCAAAATTATC | Repair template synthesis for *MOB2* S49A mutation |
| MOB2_S49A_Rev | AGAAGACGTGTATGCAGCTGAGCCTTGTGCACCTTTTGAAGATAATTTTGAtGGcGcAAA | Repair template synthesis for *MOB2* S49A mutation |
| MOB2_check_For | GGCGGGTATCCATTAAGCAA | Genotype PCR primer for detecting *MOB2* allele |
| MOB2_check_Rev | GTTTGGTCCCGCATTCATTG | Genotype PCR primer for detecting *MOB2* allele |
| sgBNI1_top | atttgAAAAGGACAACTGGGTCTGTg | CRISPR plasmid cloning against *BNI1* |
| sgBNI1_bottom | aaaacACAGACCCAGTTGTCCTTTTc | CRISPR plasmid cloning against *BNI1* |
| BNI1_S1619A_For | TTAATGAGAAAACAAATATTGGAAAGTCAACGTAAAAGaACtACaGGaTCgGTTGGCTCA | Repair template synthesis for *BNI1* S1619A mutation |
| BNI1_S1619A_Rev | TGTCAGATTCATTATTTCTTGTtGGcGcGACATTAGTTGGTGAGCCAACcGAtCCtGTaG | Repair template synthesis for *BNI1* S1619A mutation |
| BNI1_check_For | ACTGAGTCGATCCGTACCAT | Genotype PCR primer for detecting *BNI1* allele |
| BNI1_check_Rev | TTTATGTGTCGCCATCGTCA | Genotype PCR primer for detecting *BNI1* allele |
| SNR52_R_HGC1 | AGTTCATCATGTAGTACTACCAAATTAAAAATAGTTTACGCAAGTC | sgRNA synthesis against *HGC1* |
| sgRNA_F_HGC1 | GTAGTACTACATGATGAACTGTTTTAGAGCTAGAAATAGCAAGTTAAA | sgRNA synthesis against *HGC1* |
| HGC1_check_For | TGTGCGTTGTGCGTGTATAA | Genotype PCR primer for detecting *HGC1* allele |
| HGC1_check_Rev | GGTGAAAGTTAAATATGGTTGTTGT | Genotype PCR primer for detecting *HGC1* allele |
| HGC1_NAT_FLP_For | CATCAAAGCATCAAACCAATACCCAACACTTTAATATCTAGGGTTTCCATTCACATATACACATATAAACATATATTAATTAAAGGGAACAAAAGCTGGG | *HGC1* deletion construct |
| HGC1_NAT_FLP_Rev | CATCATTAAAATTTCATATCATAATAACAACATCTTTCTCCATTCTCCATTCTCTACTTTATCTTTCTCTCTTTCTTTAACTCTAGAACTAGTGGATCTG | *HGC1* deletion construct |
| SNR52_R_YCK2 | TCACATAATGTAACTGCTGGCAAATTAAAAATAGTTTACGCAAGTC | sgRNA synthesis against *YCK2* |
| sgRNA_F_YCK2 | CCAGCAGTTACATTATGTGAGTTTTAGAGCTAGAAATAGCAAGTTAAA | sgRNA synthesis against *YCK2* |
| YCK2_check_For | AATGCTACTTGGCTAATCCCA | Genotype PCR primer for detecting *YCK2* allele |
| YCK2_check_Rev | AGGAATTCCGTCACATCCTTG | Genotype PCR primer for detecting *YCK2* allele |
| YCK2_NAT_FLP_For | CGTTGAGACtTGACAACAAACCCTGCTTTGGCGGCTGCTCAAGCATCTCATAATAATATTCCTACAAAGCAAATGAATCATAAAGGGAACAAAAGCTGGG | *YCK2* deletion construct |
| YCK2_NAT_FLP_Rev | GATATCATACAACTAATGACAACACAATTTAGACCAGAACCCTTTGTTTTCTTCTTCTTCGGCAACCATTTGTTGTTGTTCTCTAGAACTAGTGGATCTG | *YCK2* deletion construct |
| HGC1_check_Rev2 | GGTGGTGATGGTCTGTTCAT | Genotype PCR primer for detecting *HGC1* allele |

**S1 Data. SNP analysis of the PR mutants**

**S2 Data. RNA-seq data**

**S3 Data. Quantitative phosphoproteomic data**

**S4 Data. Whole-proteomic data and the protein-mRNA abundance correlation**

**SUPPLEMENTAL METHODS**

**1. Protocol for transient CRISPR-Cas9 system**

The transient CRISPR system requires CaCas9 expression cassette, sgRNA expression cassette, and target gene deletion construct. The three components were all PCR amplified as described below.

**1.1. Construction of CaCas9 expression cassette**

The CaCas9 expression cassette was PCR amplified from the plasmid pV1093. The plasmid pV1093 used in this study was a kind gift from Valmik Vyas [Sci Adv 1(3):e1500248, 2015].

| Reagent | Volume (µl) |
| --- | --- |
| pV1093 (50 ng/ µl) | 1.0 |
| 10X buffer | 5.0 |
| dNTP (2.5 mM each) | 4.0 |
| 10µM CaCas9/For primer | 1.0 |
| 10µM CaCas9/Rev primer | 1.0 |
| TAKARA ExTaq | 0.25 |
| Sterilized deionized water | Up to 50.0 |

| Temperature | Time | No. of cycles |
| --- | --- | --- |
| 94 °C | 1 min |  |
| 94 °C | 30 sec | 30 cycles |
| 58 °C | 1 min |  |
| 72 °C | 4 min |  |
| 72 °C | 5 min |  |
| 4 °C | - |  |

Purify the PCR products and measure the concentration.

**1.2. Construction of sgRNA expression cassette**

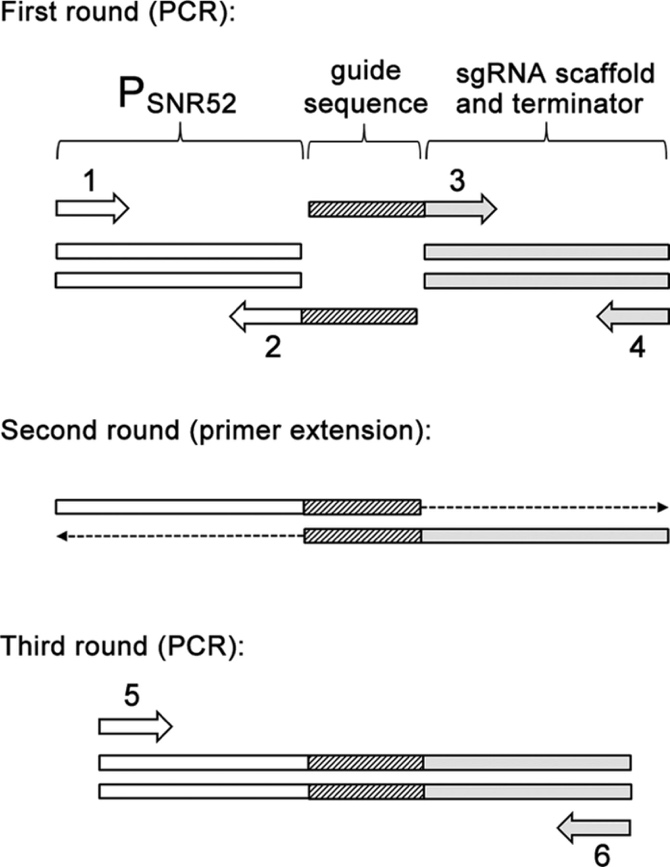

Three DNA synthesis steps fuse DNA fragments comprising the SNR52 promoter and the sgRNA scaffold. Chimeric primers 2 and 3 carry 20 complementary bases of guide sequence. The guide sequence was designed and used by Vyas et al. [Sci Adv 1(3):e1500248, 2015].

In the first step, PCR is used to create two segments of the sgRNA gene. The SNR52 promoter is amplified with primers 1 and 2; the sgRNA scaffold is amplified with primers 3 and 4. In the second step, primer extension is used to fuse the two PCR products, with chimeric extensions acting as primers. In the third step, PCR with nested primers 5 and 6 is used to amplify the final sgRNA expression cassette.

**First round PCR**

SNR52 promoter region

| Reagent | Volume (µl) |
| --- | --- |
| pV1093 (50 ng/ µl) | 1.0 |
| 10X buffer | 5.0 |
| dNTP (2.5 mM each) | 4.0 |
| 10µM primer #1 (SNR52/F) | 1.0 |
| 10µM primer #2 (SNR52/R) | 1.0 |
| TAKARA ExTaq | 0.25 |
| Sterilized deionized water | Up to 50.0 |

sgRNA scaffold region

| Reagent | Volume (µl) |
| --- | --- |
| pV1093 (50 ng/ µl) | 1.0 |
| 10X buffer | 5.0 |
| dNTP (2.5 mM each) | 4.0 |
| 10µM primer #3 (sgRNA/F) | 1.0 |
| 10µM primer #4 (sgRNA/R) | 1.0 |
| TAKARA ExTaq | 0.25 |
| Sterilized deionized water | Up to 50.0 |

| Temperature | Time | No. of cycles |
| --- | --- | --- |
| 94 °C | 1 min |  |
| 94 °C | 30 sec | 30 cycles |
| 58 °C | 1 min |  |
| 72 °C | 1 min |  |
| 72 °C | 5 min |  |
| 4 °C | - |  |

Purify the PCR products and measure the concentration.

**Second round PCR**

| Reagent | Volume (µl) |
| --- | --- |
| Purified SNR52 promoter amplicon | 2.5 |
| Purified sgRNA amplicon | 2.5 |
| 10X buffer | 2.5 |
| dNTP (2.5 mM each) | 2.0 |
| TAKARA ExTaq | 0.25 |
| Sterilized deionized water | Up to 25.0 |

*Note. Use 1: 1 molar ratio for SNR52: sgRNA amplicons. The total DNA amount of the two components should be between 100 and 1000 ng.*

| Temperature | Time | No. of cycles |
| --- | --- | --- |
| 94 °C | 2 min |  |
| 94 °C | 30 sec | 10 cycles |
| 58 °C | 10 min |  |
| 72 °C | 5 min |  |
| 72 °C | 10 min |  |
| 4 °C | - |  |

Do not need to purify the PCR products and measure the concentration.

**Third round PCR**

| Reagent | Volume (µl) |
| --- | --- |
| Second round product | 1.0 |
| 10X buffer | 5.0 |
| dNTP (2.5 mM each) | 4.0 |
| 10µM primer #5 (SNR52/N) | 1.0 |
| 10µM primer #6 (sgRNA/N) | 1.0 |
| TAKARA ExTaq | 0.25 |
| Sterilized deionized water | Up to 50.0 |

| Temperature | Time | No. of cycles |
| --- | --- | --- |
| 94 °C | 1 min |  |
| 94 °C | 30 sec | 30 cycles |
| 58 °C | 1 min |  |
| 72 °C | 1 min |  |
| 72 °C | 5 min |  |
| 4 °C | - |  |

Purify the PCR products and measure the concentration.

**1.3. Construction of gene deletion constructs**

Gene deletion constructs were synthesized by PCR using the plasmid pGR-NAT, as a template. The primers were designed to include 80 bases with homology to the sequences upstream or downstream from the target gene.

| Reagent | Volume (µl) |
| --- | --- |
| Template plasmid (50 ng/µl) | 1.0 |
| 10X buffer | 5.0 |
| dNTP (2.5 mM each) | 4.0 |
| 10µM Forward primer | 1.0 |
| 10µM Reverse primer | 1.0 |
| TAKARA ExTaq | 0.25 |
| Sterilized deionized water | Up to 50.0 |

| Temperature | Time | No. of cycles |
| --- | --- | --- |
| 94 °C | 1 min |  |
| 94 °C | 30 sec | 30 cycles |
| 58 °C | 1 min |  |
| 72 °C | 2 min |  |
| 72 °C | 5 min |  |
| 4 °C | - |  |

Purify the PCR products and measure the concentration.

**1.4. Fungal transformation**

PCR products for transformation were purified and concentrated with the commercial PCR purification kit. In the transient CRISPR system, the deletion constructs (1 ug) were co-transformed with the *CaCAS9* cassette (1 ug) and sgRNA cassette (1 ug), using the electroporation method.

**1.5. Primers**

| **Primer** | **Sequence** | **Description** |
| --- | --- | --- |
| CaCas9/for | ATCTCATTAGATTTGGAACTTGTGGGTT | Forward and reverse primers for amplification of *CaCas9* cassette |
| CaCas9/rev | TTCGAGCGTCCCAAAACCTTCT |  |
| SNR52/F | AAGAAAGAAAGAAAACCAGGAGTGAA | Forward primer for amplification of *SNR52* promoter |
| SNR52/R | (Reverse complementary of 20-nt target) + CAAATTAAAAATAGTTTACGCAAGTC | Reverse primer for amplification of *SNR52* promoter with overlapping guide sequence |
| sgRNA/F | (20-nt target sequence) + GTTTTAGAGCTAGAAATAGCAAGTTAAA | Forward primer for amplification of sgRNA scaffold with overlapping guide sequence |
| sgRNA/R | ACAAATATTTAAACTCGGGACCTGG | Reverse primer for amplification of sgRNA scaffold |
| SNR52/N | GCGGCCGCAAGTGATTAGACT | Forward and reverse nested primers for third round PCR for construction of sgRNA expression cassette |
| sgRNA/N | GCAGCTCAGTGATTAAGAGTAAAGATGG |  |

**2. Large CRISPR-mediated deletion**

**KM11** (*bcy1∆*/*BCY1 cyr1∆*/*∆*): We first made the KM11 strain without using the CRISPR system. The parental *cyr1∆*/*∆* strain was made in the previous study (Parrino et al., 2017) and the NAT selection marker (SAT1-FLP) was excised for marker recycling. The *BCY1* deletion construct was synthesized with the BCY1_NAT_FLP_For and BCY1_NAT_FLP_Rev primers, using the plasmid pGR-NAT plasmid as a template. The construct was transformed into the *cyr1∆*/*∆* cells and the heterozygous *BCY1* deletion mutants were selected for NAT resistance. PCR genotyping of the transformants verified the heterozygous deletion. The NAT selection marker (SAT1-FLP) was excised again for large CRISPR-mediated deletion.

**KM12** (*Chr2L 270kb deletion*): When deleting 270kb region of chromosome 2 for gene mapping, two sgRNA expression cassettes were used to cut the C2_00030W and C2_01540W loci. Chimeric primers SNR52_R_30 and sgRNA_F_30 were used to synthesize the sgRNA cassette against C2_00030W. Chimeric primers SNR52_R_1540 and sgRNA_F_1540 were used to synthesize the sgRNA cassette against C2_01540W. The 270kb deletion construct was synthesized with the 30_NAT_FLP_For and 1540_NAT_FLP_Rev primers, using the plasmid pGR-NAT plasmid as a template. The *CaCAS9* expression cassette, two sgRNA expression cassettes, and deletion construct were co-transformed into the Nat^s^ KM11 strain. Nat^r^ transformants were selected, and PCR genotyping of the transformants verified the heterozygous deletion.

**KM13** (*Chr2L 590kb deletion*): When deleting 590kb region of chromosome 2 for gene mapping, two sgRNA expression cassettes were used to cut the C2_00030W and C2_02960C loci. Chimeric primers SNR52_R_30 and sgRNA_F_30 were used to synthesize the sgRNA cassette against C2_00030W. Chimeric primers SNR52_R_2960 and sgRNA_F_2960 were used to synthesize the sgRNA cassette against C2_02960C. The 590kb deletion construct was synthesized with the 30_NAT_FLP_For and 2960_NAT_FLP_Rev primers, using the plasmid pGR-NAT plasmid as a template. The *CaCAS9* expression cassette, two sgRNA expression cassettes, and deletion construct were co-transformed into the Nat^s^ KM11 strain. Nat^r^ transformants were selected, and PCR genotyping of the transformants verified the heterozygous deletion.

**KM14** (*Chr2L 90kb deletion*): When deleting 90kb region of chromosome 2 for gene mapping, two sgRNA expression cassettes were used to cut the C2_00030W and C2_00550W loci. Chimeric primers SNR52_R_30 and sgRNA_F_30 were used to synthesize the sgRNA cassette against C2_00030W. Chimeric primers SNR52_R_550 and sgRNA_F_550 were used to synthesize the sgRNA cassette against C2_00550W. The 90kb deletion construct was synthesized with the 30_NAT_FLP_For and 550_NAT_FLP_Rev primers, using the plasmid pGR-NAT plasmid as a template. The *CaCAS9* expression cassette, two sgRNA expression cassettes, and deletion construct were co-transformed into the Nat^s^ KM11 strain. Nat^r^ transformants were selected, and PCR genotyping of the transformants verified the heterozygous deletion.

**KM15** (*Chr2L 90kb→180kb deletion*): When deleting 90kb→180kb region of chromosome 2 for gene mapping, two sgRNA expression cassettes were used to cut the C2_00560W and C2_01140C loci. Chimeric primers SNR52_R_560 and sgRNA_F_560 were used to synthesize the sgRNA cassette against C2_00560W. Chimeric primers SNR52_R_1140 and sgRNA_F_1140 were used to synthesize the sgRNA cassette against C2_01140C. The 90kb→180kb deletion construct was synthesized with the 560_NAT_FLP_For and 1140_NAT_FLP_Rev primers, using the plasmid pGR-NAT plasmid as a template. The *CaCAS9* expression cassette, two sgRNA expression cassettes, and deletion construct were co-transformed into the Nat^s^ KM11 strain. Nat^r^ transformants were selected, and PCR genotyping of the transformants verified the heterozygous deletion.

**KM16** (*Chr2L 180kb→270kb deletion*): When deleting 180kb→270kb region of chromosome 2 for gene mapping, two sgRNA expression cassettes were used to cut the C2_01150W and C2_01540W loci. Chimeric primers SNR52_R_1150 and sgRNA_F_1150 were used to synthesize the sgRNA cassette against C2_1150W. Chimeric primers SNR52_R_1540 and sgRNA_F_1540 were used to synthesize the sgRNA cassette against C2_01540W. The 590kb deletion construct was synthesized with the 1150_NAT_FLP_For and 1540_NAT_FLP_Rev primers, using the plasmid pGR-NAT plasmid as a template. The *CaCAS9* expression cassette, two sgRNA expression cassettes, and deletion construct were co-transformed into the Nat^s^ KM11 strain. Nat^r^ transformants were selected, and PCR genotyping of the transformants verified the heterozygous deletion.

**KM17** (*Chr2L 180kb deletion*): When deleting 180kb region of chromosome 2 for gene mapping, two sgRNA expression cassettes were used to cut the C2_00030W and C2_01140C loci. Chimeric primers SNR52_R_30 and sgRNA_F_30 were used to synthesize the sgRNA cassette against C2_00030W. Chimeric primers SNR52_R_1140 and sgRNA_F_1140 were used to synthesize the sgRNA cassette against C2_01140C. The 180kb deletion construct was synthesized with the 30_NAT_FLP_For and 1140_NAT_FLP_Rev primers, using the plasmid pGR-NAT plasmid as a template. The *CaCAS9* expression cassette, two sgRNA expression cassettes, and deletion construct were co-transformed into the Nat^s^ KM11 strain. Nat^r^ transformants were selected, and PCR genotyping of the transformants verified the heterozygous deletion.

**KM18** (*Chr2L 90kb→270kb deletion*): When deleting 90kb→270kb region of chromosome 2 for gene mapping, two sgRNA expression cassettes were used to cut the C2_00560W and C2_01540W loci. Chimeric primers SNR52_R_560 and sgRNA_F_560 were used to synthesize the sgRNA cassette against C2_00560W. Chimeric primers SNR52_R_1540 and sgRNA_F_1540 were used to synthesize the sgRNA cassette against C2_01540W. The 90kb→270kb deletion construct was synthesized with the 560_NAT_FLP_For and 1540_NAT_FLP_Rev primers, using the plasmid pGR-NAT plasmid as a template. The *CaCAS9* expression cassette, two sgRNA expression cassettes, and deletion construct were co-transformed into the Nat^s^ KM11 strain. Nat^r^ transformants were selected, and PCR genotyping of the transformants verified the heterozygous deletion.

**KM19** (*Chr2L 90kb→260kb deletion*): When deleting 90kb→260kb region of chromosome 2 for gene mapping, two sgRNA expression cassettes were used to cut the C2_00560W and C2_01500W loci. Chimeric primers SNR52_R_560 and sgRNA_F_560 were used to synthesize the sgRNA cassette against C2_00560W. Chimeric primers SNR52_R_1500 and sgRNA_F_1500 were used to synthesize the sgRNA cassette against C2_01500W. The 90kb→260kb deletion construct was synthesized with the 560_NAT_FLP_For and 1500_NAT_FLP_Rev primers, using the plasmid pGR-NAT plasmid as a template. The *CaCAS9* expression cassette, two sgRNA expression cassettes, and deletion construct were co-transformed into the Nat^s^ KM11 strain. Nat^r^ transformants were selected, and PCR genotyping of the transformants verified the heterozygous deletion.

**KM20** (*Chr2L 90kb→250kb deletion*): When deleting 90kb→250kb region of chromosome 2 for gene mapping, two sgRNA expression cassettes were used to cut the C2_00560W and C2_01460C loci. Chimeric primers SNR52_R_560 and sgRNA_F_560 were used to synthesize the sgRNA cassette against C2_00560W. Chimeric primers SNR52_R_1460 and sgRNA_F_1460 were used to synthesize the sgRNA cassette against C2_01460C. The 90kb→250kb deletion construct was synthesized with the 560_NAT_FLP_For and 1460_NAT_FLP_Rev primers, using the plasmid pGR-NAT plasmid as a template. The *CaCAS9* expression cassette, two sgRNA expression cassettes, and deletion construct were co-transformed into the Nat^s^ KM11 strain. Nat^r^ transformants were selected, and PCR genotyping of the transformants verified the heterozygous deletion.
